# *INTERMEDIUM-C* mediates the shade-induced bud growth arrest in barley

**DOI:** 10.1101/2020.07.30.228510

**Authors:** Hongwen Wang, Christiane Seiler, Nese Sreenivasulu, Nicolaus von Wirén, Markus Kuhlmann

**Affiliations:** Department of Physiology and Cell Biology, Leibniz Institute of Plant Genetics and Crop Plant Research (IPK) Gatersleben, Corrensstrasse 3, 06466 Stadt Seeland, Germany; Department of Molecular Genetics, Leibniz Institute of Plant Genetics and Crop Plant Research (IPK) Gatersleben, Corrensstrasse 3, 06466 Stadt Seeland, Germany; International Rice Research Institute (IRRI), Grain Quality and Nutrition Center, Metro Manila, Philippines

**Author notes:** Correspondence to: Hongwen Wang, Nicolaus von Wirén and Markus Kuhlmann, Department of Molecular Genetics & Department of Physiology and Cell Biology, Leibniz Institute of Plant Genetics and Crop Plant Research (IPK), Gatersleben, Corrensstrasse 3, 06466 Stadt Seeland, Germany. The authors responsible for the distribution of materials integral to the findings presented in this article in accordance with the policy described in the instructions for authors are Hongwen Wang, Nicolaus von Wirén and Markus Kuhlmann.

**Keywords:** *INTERMEDIUM-C*, yield, barley, bud growth arrest, shade avoidance, decapitation, abscisic acid

## Abstract

Tiller formation is a key agronomic determinant for grain yield in cereal crops. The modulation of this trait is controlled by transcriptional regulators and plant hormones, tightly regulated by external environmental conditions. While endogenous (genetics) and exogenous (environmental factors) triggers for tiller formation have mostly been investigated separately, it has remained elusive how they are integrated into the developmental program of this trait. The transcription factor *INTERMEDIUM-C (INT-C)*, which is the barley ortholog of the maize domestication gene *TEOSINTE BRANCHED1 (TB1)* has a prominent role in regulating tiller bud outgrowth. Here we show that *INT-C* is expressed in tiller buds, required for bud growth arrest in response to shade. In contrast to wild type plants, *int-c* mutant plants are impaired in their shade response and do not stop tiller production after shading. Gene expression levels of *INT-C* are up-regulated under light-limiting growth conditions, and down-regulated after decapitation. Transcriptome analysis of wild-type and *int-c* buds under control and shading conditions identified target genes of INT-C that belong to auxin and gibberellin biosynthesis and signaling pathways. Our study identifies INT-C as integrator of the shade response into tiller formation, which is prerequisite for implementing shading responses in the breeding of cereal crops.

## Introduction

Ensuring yield stability of cereal crops is a major requirement for plant breeding in the face of climate change (Kang et al., 2009). In particular, breeding of new elite varieties with optimized shoot architecture, such as tiller number, will allow maintaining high yield potential in unfavorable environments. In many countries, particularly at very high latitudes or on shaded slopes, light is a limiting factor affecting crop growth and yield. Moreover, during the past decades intensive crop management has increased sowing and plant stand densities to improve homogeneity among individual plants that now produce less tillers. To increase light capture, plants have evolved refined mechanisms to maximize light harvesting and/or to prevent shade, i.e. low light intensity and/or a low ratio of red (R) to far-red (FR) light. Suboptimal light triggers a suite of phenotypic changes, defined as the shade avoidance response (SAR) that includes hypocotyl and petiole elongation, an upward orientation of leaves and early flowering. Additionally, a common characteristic of SAR is the suppression of shoot branching in a wide variety of species (Smith and Jordan, 1994; Tucic et al., 2006; Aguilar-Martinez et al., 2007; Gonzalez-Grandio et al., 2013).

The extensive work carried out in Arabidopsis has provided a detailed understanding of the SAR (Wang and Wang, 2015). By sensing and responding to R and FR light, the five photoreceptors of the phytochrome family (phyA-phyE) regulate a variety of developmental processes (Franklin and Quail, 2010). Among these, phytochrome B (phyB) appears to be the most important photoreceptor involved in shade detection and avoidance (Reed et al., 1993; Ballare, 1999), functioning redundantly with other members of its clade (Stamm and Kumar, 2010). Phytochrome proteins act as dimers and exist in two photo-convertible forms: ‘Pr’ (the red-light-absorbing, inactive form) and ‘Pfr’ (the far-red-light-absorbing, active form), with the Pfr:Pr ratio reflecting the R:FR ratio of the environment (Smith, 2000). Upon photo-conversion into active Pfr, phytochromes migrate to the nucleus, where they regulate gene expression by interacting with several basic helix-loop-helix (bHLH) transcription factors, including phytochrome interacting factor (PIF) and PIF-like (PIL) proteins (Chen et al., 2004; Duek and Fankhauser, 2005). In many plant species, genome-wide transcriptional dynamics of the SAR have been studied in petioles or leaf blades at the seedling stage, while SAR is less defined in economically important monocots (Devlin et al., 2003; Tao et al., 2008; Hornitschek et al., 2012; Wang et al., 2016).

In the agricultural production of graminaceous crop species, plant density is a major determinant for crop yield. Shading caused by elevated plant densities reduces not only photosynthetic active radiation (PAR) flux density but also the R/FR ratio of the light reaching the lower strata of the canopy. Generally, increasing plant density results in progressively stronger suppression of tillering due to accelerated apical shoot development and stem elongation. This pattern continues until the beginning of the stem elongation phase (Zadoks, 1985). Moreover, early-emerging tillers contribute more to grain yield than do tillers that emerge later. The regulation of barley tiller outgrowth by shade has also been supported by the observation that supplemental FR illumination of elongating leaves or of the main stem base reduced the total number of tillers per plant (Skinner and Simmons, 1993). In spite of its great ecological and economic impact, little is known about the underlying molecular mechanisms linking shade-initiated transcriptional changes with the suppression of tiller outgrowth in barley plants, especially regarding the tiller bud that is one of the most important sites of shade action.

Lateral shoot growth is coordinately controlled by conserved interactions that regulate the biosynthesis and signaling of the hormones auxin, abscisic acid (ABA), strigolactones (SL), and cytokinins (CK). Auxin and SL, synthesized mainly in the shoot apex and root, respectively, inhibit branching, while CK, synthesized mostly in the root and stem, promote branching (Kebrom et al., 2013; Wang et al., 2018).

The class II TEOSINTE BRANCHED1, CYCLOIDEA, and PCF (TCP) transcription factors TEOSINTE BRANCHED 1 (TB1)-like, in monocots, and BRANCHED1 (BRC1)-like, in dicots, act locally inside the axillary bud where they are subject to transcriptional regulation by hormonal cross-talk (auxin-SL-CK) and cause bud growth arrest (Doebley et al., 1997; Aguilar-Martinez et al., 2007; Minakuchi et al., 2010; Martin-Trillo et al., 2011; Braun et al., 2012). Moreover, recent studies have implicated the involvement of TB1 orthologs in the shade-induced response of branch suppression. In *Sorghum bicolor* and maize, active phyB (Pfr) suppresses the expression of the *TB1* gene and induces bud outgrowth. On the other hand, light signals that inactivate phyB allow increased expression of *TB1* and suppression of bud outgrowth (Kebrom et al., 2006; Kebrom et al., 2010; Whipple et al., 2011). In Arabidopsis, *BRC1* is upregulated in axillary buds of plants grown at high density and is required for complete branch suppression in these conditions (Aguilar-Martinez et al., 2007). These results suggest that the phytochrome pathway is involved in the control of *TB1* and axillary meristem outgrowth, and provides a link between environmental variation and gene action controlling branching. In barley, *INT-C* is an ortholog of the maize domestication gene *TB1*, and its mutants exhibit higher tiller numbers and enlarged lateral spikelets (Ramsay et al., 2011).

Here, we investigated the shade avoidance response of early formed tiller buds in barley plants before the rapid stem elongation stage, and studied the role of *INT-C* during the SAR by comparable transcriptome analysis. The analysis revealed that the dynamics of *INT-C* expression during SAR is critical for genome-wide reprogramming of gene expression and that the gene categories affected support a central role of *INT-C* in tiller bud arrest. A comparison of genes responding to shade-induced bud arrest and decapitation-triggered bud activation allowed us to identify key regulators influencing *bud dormancy* and *bud activation* genes. Thus, these findings would enable us to better understand the genetic mechanisms controlling the reversible transition of growth to dormancy in barley tiller buds.

## Materials and Methods

### Plant material and growth conditions

*Hordeum vulgare* cv. Bowman, a two-rowed spring barley cultivar, was used as wild type for comparison to its near isogenic mutant BW421 (*int-c*.*5*) (Ramsay et al., 2011). For the experimental analyses except the shading experiment, wild type and *int-c* mutant plants were cultivated in a greenhouse. Seeds were sown in either 54 or 96 well trays and germinated in a climate chamber for 10 days at 11°C day and 7°C night temperature under 16 h light. After that, seedlings were transferred to pots (diameter 16 cm) filled with 2 parts of compost, 2 parts of “Substrat 2” (Klasmann) and 1 part of quartz sand, and allowed to grow in the greenhouse at 20°C day/14°C night under 16 h light. For the density experiments, three barley planting densities (1, 5, 10 plants pot^-1^) were established in pots (size: 21-liter, top diameter: 310 cm, base diameter: 250 cm, height: 214 cm), and the planting density test was repeated four times. For the decapitation experiments, when plants were grown till the early stem elongation stage, the shoot apices were removed, and the apical dominance test was repeated three times. For the shading experiments, seedlings were grown in a climate-controlled growth chamber at a temperature of 12°C, 70% humidity and a 12/12 h day/night cycle. Green shading was achieved by using a green plastic filter (122 Fern Green; LEE Filters, Andover, UK) (Kegge et al., 2013).

### Sequence retrieval for TCP proteins

In order to identify barley genes putatively encoding TCP transcription factors, the latest barley annotation and genome sequence at http://webblast.ipk-gatersleben.de/barley/viroblast.php was searched using the TBLASTN algorithm with Arabidopsis TCP proteins or TCP domains as query sequence. All redundant sequences were discarded from further analysis based on ClusterW (Thompson et al., 1994). Furthermore, to verify the reliability of the initial results, all non-redundant candidate TCP sequences were analyzed to confirm the presence of the conserved TCP domain using the InterproScan program (Quevillon et al., 2005). The sequences of TCP family members in the genome of Arabidopsis were retrieved from the PlantTFDB plant transcription factor database (http://planttfdb.cbi.pku.edu.cn/, v3.0). As for *Antirrhinum majus* and *Oryza sativa* TCP sequences, they were obtained from NCBI.

### Phylogenetic analysis

Multiple sequence alignments were conducted on the amino acid sequences of TCP proteins in Arabidopsis and barley genomes using Cluster X (Thompson et al., 1997) with default settings. Subsequently, MEGA 6.0 software (Tamura et al., 2013) was employed to construct an unrooted phylogenetic tree based on alignments using the Neighbor-Joining (NJ) method with the following parameters: JTT model, pairwise gap deletion and 1,000 bootstraps.

### Extraction and quantification of ABA

ABA was extracted from fresh plant materials using ethyl acetate (100%). Isotopically labelled D6-ABA was used as an internal standard and added to each sample during the extraction procedure. Extraction was carried out twice with 1 ml of ethyl acetate at 4°C. The supernatant collected after centrifugation (13.000 g, 10 min, and 4°C) was evaporated to dryness at room temperature using a vacuum concentrator. The dried samples were dissolved in acetonitrile: methanol (1:1) and filtered using a 0.8 µm filter (Vivaclear). The filtrate (10 µL) was used for subsequent quantification by LC-MS/MS (Dionex Summit coupled to Varian 1200 L). Chromatographic separation was carried out on a C18 column (4 µm, 100 mm; GENESIS; Vydac/USA). MRM and quantification was done using the mass traces 263/153 for ABA and 269/159 for D6-ABA. The validity of the extraction and measurement procedure was verified in recovery experiments (approx. 82-95%). Quantification was based on calibration with known ABA standards and individual recovery rates for the samples, as described in (Kong et al., 2008).

### qRT-PCR and microarray hybridization & data analyses

RNA was extracted from fresh plant tissues from independent or pooled biological replicates with the same treatment using a plant mini RNA kit (Qiagen, Hilden, Germany) following the manufacturer’s protocol, and its quality and quantity assessed with a Nano drop device (Peqlab, Erlangen, Germany). A 500-ng aliquot was taken as template for the synthesis of the first cDNA strand, primed by oligo(dT), using a RevertAid cDNA kit (ThermoFisher SCIENTIFIC, Waltham, MA, USA). The subsequent qRT-PCR was based on the Power SYBR^®^ Green PCR Master Mix (ThermoFisher SCIENTIFIC) and conducted in an Applied Biosystems 7900HT Fast Real-Time PCR system (ThermoFisher SCIENTIFIC) following the manufacturer’s protocol. Relative transcript abundances were obtained using _ΔΔ_C_T_ method (Livak and Schmittgen, 2001) and were normalized against the abundance of serine/threonine phosphatase PP2A transcript. The primer sequences employed are given in Supplemental Table S6. The presence of a unique PCR product was verified by dissociation analysis and each qRT-PCR was repeated at least three times. Each biological replicate was represented by three technical replicates.

For the microarray procedure, the same RNA samples extracted from three biological replicates was used and the quality of the RNA was verified with a Bioanalyzer 2100 device (Agilent Technologies, Santa Clara, CA, USA). The RNA was labeled by using the Low input QuickAmp Labeling kit (Agilent Technologies) and hybridized, following the manufacturer’s protocol, to a custom-synthesized 60 k Barley Microarray (Agilent Technologies) (Koppolu et al., 2013). The resulting data were analysed using GeneSpring 13.0 GX software (Agilent Technologies). After quantile normalization and baseline transformation to the median of all samples, the probesets (genes) were filtered by Coefficient of Variation <50%, followed by moderated T-Test and Bonferroni-Holm multiple testing corrections. Probesets passing the P-value cut-off <0.05 with a fold change of ≥2.0 were selected as differentially expressed genes (DEG). Analyses of functional categories with *INT-C*-dependent upregulated and downregulated genes were performed using MapMan. The fold enrichment was calculated as follows: (number in class input_set/number of total input_set)/ (number in class reference_set/ number of total reference_set). The *P* value was determined by a hypergeometric distribution test (R Core Team, 2013). The data was sorted by fold enrichment with a cut-off of *P* < 0.05. For the specific pathway enrichment analysis by Wilcoxon-Rank-Sum test was implemented in MapMan. The resulting enrichments for the functional classes (p < 0.05, determined by a hypergeometric test; R Core Team, 2013) provided a map of gene modules regulated in dependence of *INT-C* during SAR. To confirm that the common genes of the two identified groups are not by random, the test of statistical significance was applied by a web-based tool at http://nemates.org/MA/progs/overlap_stats.html. The co-regulated genes were retrieved from Genevestigator (Zimmermann et al., 2004), and Gene Ontology analysis was performed in agriGO v2.0 with default parameters (Tian et al., 2017).

### Statistical analysis

To test the statistical significance of the data, Student’s *t* test and one-way analysis of variance (ANOVA) with Tukey test for significance were used. Asterisks denote significant differences in Student’s *t* tests. Different letters denote significant differences in Tukey’s test.

## Results

### *INT-C* is a member of the barley TCP gene family

*<u>INT</u>ERMEDIUM-<u>C</u>* (*INT-C*) is the barley orthologue of *TEOSINTE BRANCHED1* (Ramsay et al., 2011), one of the name giving members of the well-studied <u>T</u>EOSINTE BRANCHED1, <u>C</u>YCLOIDEA, <u>P</u>ROLIFERATING CELL NUCLEAR ANTIGEN BINDING FACTOR (TCP) gene family (Li, 2015). TCP transcription factors are defined by the TCP domain, which is a 59 aa-long <u>b</u>asic <u>h</u>elix-loop-<u>h</u>elix (bHLH) structure which provides the ability to bind GC-rich DNA sequence motifs (Martin-Trillo and Cubas, 2010). In barley, INT-C is encoded by *HORVU4Hr1G007040* and was identified as corresponding gene of the *vrs5* locus on chromosome 4 (Ramsay et al., 2011) being involved in the fertility of lateral spikelets.

To identify the genes closest to *INT-C*, the barley TCP gene family was analyzed using the latest barley annotation and genome sequence at http://webblast.ipk-gatersleben.de/barley/viroblast.php and the TBLASTN algorithm with Arabidopsis TCP proteins or TCP domains as query sequence (Figure 1A; Supplemental Table S1). This family consists of 20 genes which contain a putative TCP-helix-loop-helix-type domain at the N-terminus (Figure 1B). The phylogenetic tree which was built based on multiple alignments of the TCP domain in TCP proteins showed that barley TCP proteins could be divided into two groups, as for all species so far (Figure 1A/B). The class I group was formed by nine predicted proteins related to the PCF rice factors (Kosugi and Ohashi, 1997), while class II was comprised of eleven predicted proteins related to the *Antirrhinum CYC* and *CIN* genes and to *OsTB1* (Luo et al., 1996; Doebley et al., 1997; Nath et al., 2003; Takeda et al., 2003). In addition, the class II group could be further divided into two subclades: the CIN group formed by eight members and the CYC/TB1 group formed by three members. *HvTC16* and *HvTC17* from the class II CYC/TB1 contain an R domain that is also found in *HvTC12* from the class II CIN group (Figure 1C), as previously described in Arabidopsis (Yao et al., 2007).

**Figure 1.**
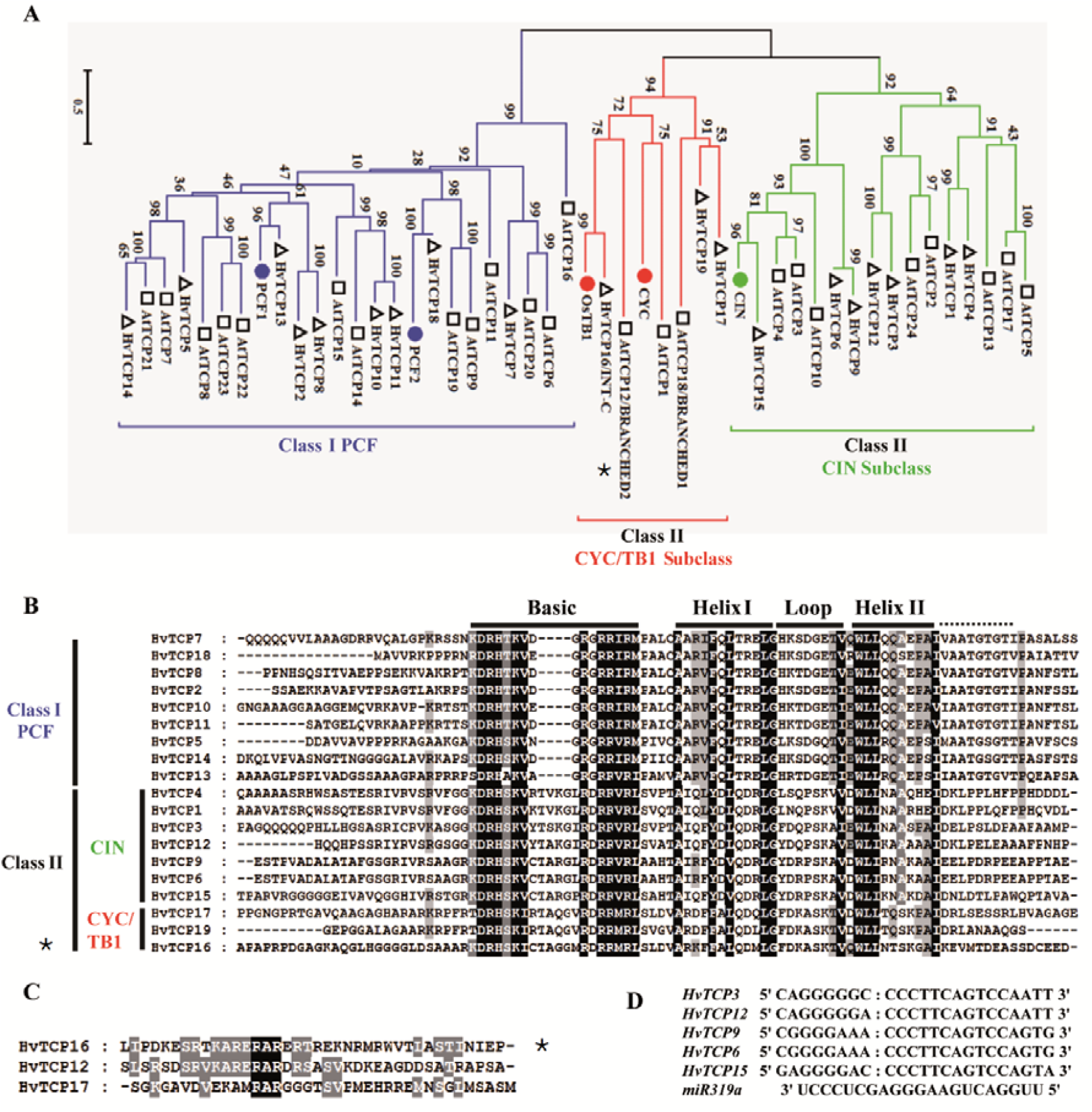
Phylogenetic relations of Arabidopsis and barley TCP proteins. **A)** The phylogenetic tree of TCP proteins was built based on multiple alignment of the TCP domain amino acid sequence using the Neighbor-Joining method with 1000 bootstrap replicates. Blue, red and green lines indicate the PCF, CYC/TB1 and CIN clades, respectively. Each Arabidopsis protein is indicated by a square, each barley protein is indicated by a triangle. **B)** Alignment of the TCP domain and adjoining sequence for the predicted barley TCP proteins. Overall conserved amino acids are shaded in black. Amino acids 80% or 100% conserved in Class II or Class I are shaded in light gray and dark gray, respectively. The Basic, Helix I, Loop, and Helix II regions are indicated. **C)** Alignment of the R-domain of Class II subfamily members. Amino acids are expressed in the standard single letter code. Sequences were aligned with ClustalW and represented with Genedoc. **D)** Alignment of putative target areas for *miR319* (aligned in reverse). *INT-C* (*HvTCP16)*: asterisks

Although in eudicots several CYC/TB1 sequences are found, and phylogenetic analyses have suggested that duplications within this clade occurred at the base of eudicots, in monocots only one type of CYC/TB1 has been identified (e.g., Os*TB1*) (Howarth and Donoghue, 2006). Our phylogenetic analysis revealed that, based on the absence of the R domain, none of the newly identified barley TCP genes can be considered as paralogue of *INT-C*.

As described for the model plant Arabidopsis, five of the CIN subclade members are post-transcriptionally regulated by *miRNA319* (*AtTCP2, 3, 4, 10* and *24*) (Palatnik et al., 2003; Ori et al., 2007; Palatnik et al., 2007). In barley *hvu-miR319a* (UUGGACUGAAGGGAGCUCCC) is encoded by CL16998_Contig1 (Ozhuner et al., 2013) and expressed in roots, while it is not detectable in leaf tissue. The closest barley homologs of these Arabidopsis genes are the five genes, *HvTCP3, HvTCP6, HvTCP9, HvTCP12*, and *HvTCP15*. These barley CIN subclade members contain sequences with putative binding sites for *hvu-miR319* (Supplemental Table S1). Figure 1D shows the alignment of the target sites of these genes with the *miR319* sequence. This suggests that regulation of leaf development by a redundant set of miRNA-regulated homologous TCP genes occurs in barley, while *INT-C* does not represent a miR319 target.

### *INT-C* loss-of-function leads to an early promotion of tiller bud outgrowth in barley

Previous studies noticed that *int-c* mutants in various cultivars produce more tillers than the respective wild type plants during early vegetative stages (Ramsay et al., 2011; Liller et al., 2015). However, this increased tiller formation did not translate into more productive tillers. In contrast the number of tillers was significantly lower at the point of harvest. The growth experiment described here performed with the near isogenic mutant BW421 (*int-c*.*5*) and the corresponding wild type confirmed these earlier observations. An increase in tiller number in *int-c* was only detectable at the early developmental stages (Figure 2A) between two and five weeks after germination. At later developmental stages (6 to 8 weeks after germination), the pattern of tiller number was reversed. This observation correlated with an earlier anthesis of *int-c* compared to the wild type cv. Bowman (Supplementary Figure 1), leading to an earlier arrest of tiller bud production in *int-c*. The tillers of barley are formed in a sequential order, starting with the first tiller bud under the coleoptile. The development of the tiller buds in the axils of successive leaves was studied by dissecting the plants at different developmental stages (Figure 2B). To investigate the involvement of *INT-C* in bud initiation and bud outgrowth, the primary tiller buds were classified (Figure 2C/D) as dormant bud (800∼1200 μm), outgrowing bud (1.5∼100 mm) or tiller emergence (10∼35 cm) in each leaf axil at an early developmental stage (2 to 3 weeks after germination). Figure 2D indicates the enhanced bud outgrowth in *int-c* mutants compared to the wild type. This result suggests that the outgrowth of tiller buds is accelerated in *int-c* mutants at early developmental stages but is slowed down at later stages (> 5 weeks).

**Figure 2.**
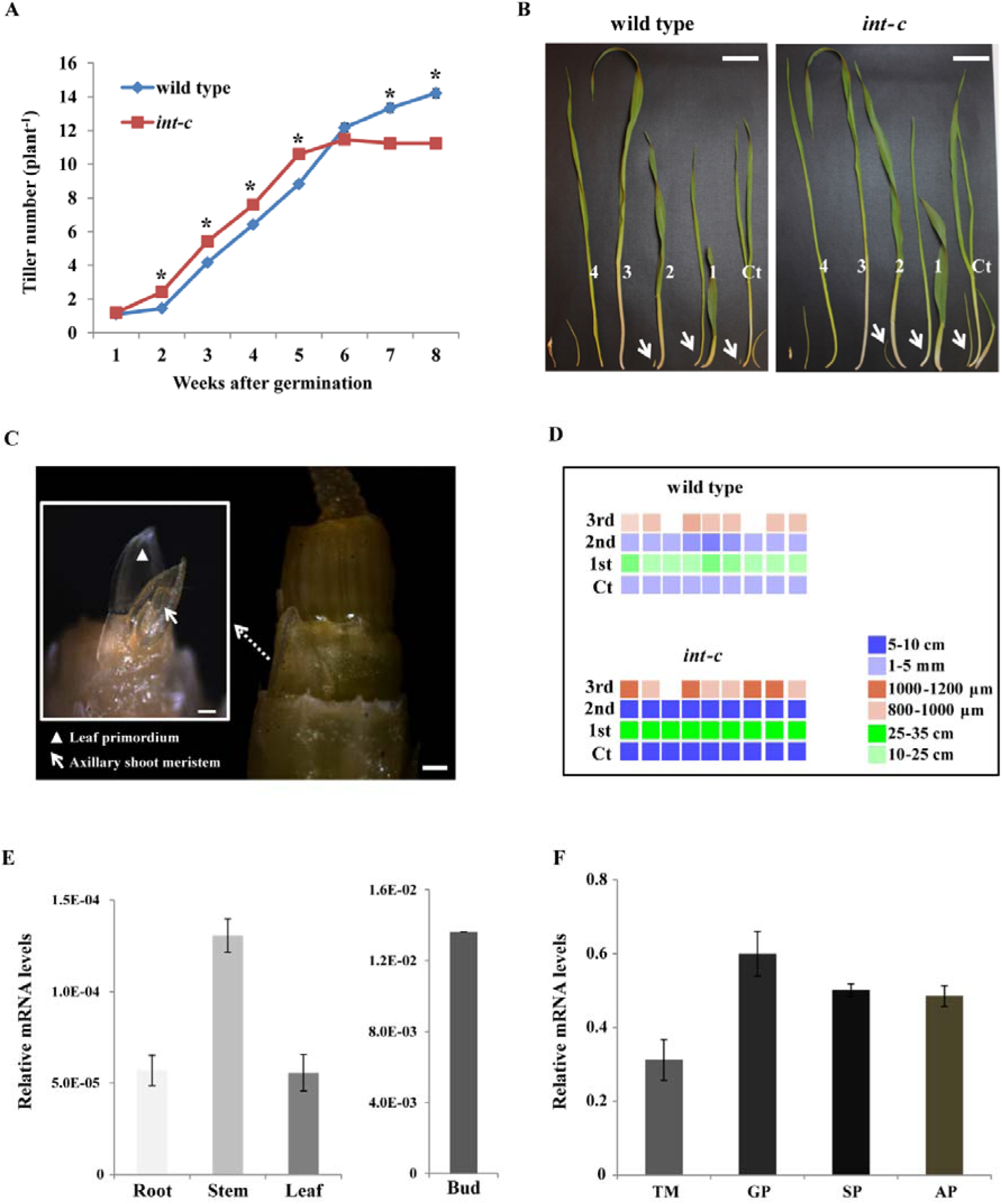
*INT-C* is involved in barley plant architecture by tiller bud outgrowth. Analysis of tiller development in wild type and *int-c* plants. **A)** Tiller number of *int-c* and wild type plants 1-8 weeks after germination (*n* = 25-30 plants). Asterisks indicate significant differences (Student’s *t* test, P < 0.001) between wild type and *int-c* mutant plants. **B)** Dissected tillers from successive leaf axils in about 2 to 3-week-old seedlings. Ct, coleoptile tiller; 1-4, order of leaves; bars = 5 cm; arrows, tiller buds. **C)** Exemplary tiller bud formation stage in the third leaf axil. The area of the close-up view is outlined with a white box in the left image. Dissection of a tiller bud at this stage will reveal a shoot apex with leaf primordia and meristematic dome. Scale bars represent 200 μm. **D)** Schematic representations of tiller bud production in each leaf axil of the wild type and *int-c* in 2- to 3-week-old seedlings. Each column stands for a single plant, and each row stands for a leaf axil in order from bottom to top, starting with the coleoptile tiller. Different color squares denote different tiller bud lengths. **E)** INT-C (HvTCP16) mRNA levels in different tissues and F) during spike development as analyzed by real-time qRT-PCR. Bars represent means ± SD; n = 3 biological replicates. Serine/threonine protein phosphatase HvPP2A-4 mRNA was used as a reference. TM, triple mound; GP, glume primordium; SP, stamen primordium; AP, awn primordium.

*INT-C* mRNA levels were analyzed by real-time qRT-PCR in different tissues and during spike development (Figure 2E/F). *INT-C* mRNA was detectable at highest levels in tiller buds, supporting its role in the control of tiller bud development. It was expressed at lower levels in other tissues such as root, stem and leaf. During spike meristem development mRNA levels of *INT-C* peaked at the glume primordium stage. In the later stages (stamen primordium and awn primordium) relative high levels of *INT-C* mRNA were persisting. This peak of expression correlated with the observation that after awn primordium stage profound differences in the development of lateral spikelet in *int-c* occurred compared to wild type (Ramsay et al., 2011).

### *INT-C* mRNA abundance decreased after decapitation

Apical dominance is the inhibitory control exerted by the shoot apex over the outgrowth of the lateral buds (Cline, 1997). Decapitation stimulates bud reactivation after breaking the apical dominance (Hall and Hillman, 1975; Napoli et al., 1999; Cline, 2000; Tatematsu et al., 2005; Aguilar-Martinez et al., 2007). To analyze the involvement of *INT-C* in integrating the decapitation response into bud outgrowth we compared barley *int-c* mutant plants to the respective wild type plants. For this approach, three-week-old plants that had undergone early stem elongation, were decapitated. Two weeks later, two tiller buds of decapitated Bowman plants had elongated prematurely. Thus, total tiller number in decapitated wild-type plants reached the same level as in *int-c* mutants, in which decapitation had no effect (Figure 3A). To investigate whether this response was related with a transcriptional downregulation of *INT-C*, mRNA levels in axillary buds were analyzed by real time qRT-PCR after decapitation, before any visible sign of bud outgrowth (Figure 3B). *INT-C* mRNA decreased significantly 6 h after decapitation. *DRM1*/*ARP* (*DORMANCY-ASSOCIATED GENE*/*AUXIN-REPRESSED PROTEIN*) mRNA, an early marker for bud dormancy (Stafstrom et al., 1998; Tatematsu et al., 2005), also showed reduction and reached its minimum 24 h after decapitation. These results support the idea that *INT-C* is involved in the early response to bud release from apical dominance and required for bud activation.

**Figure 3.**
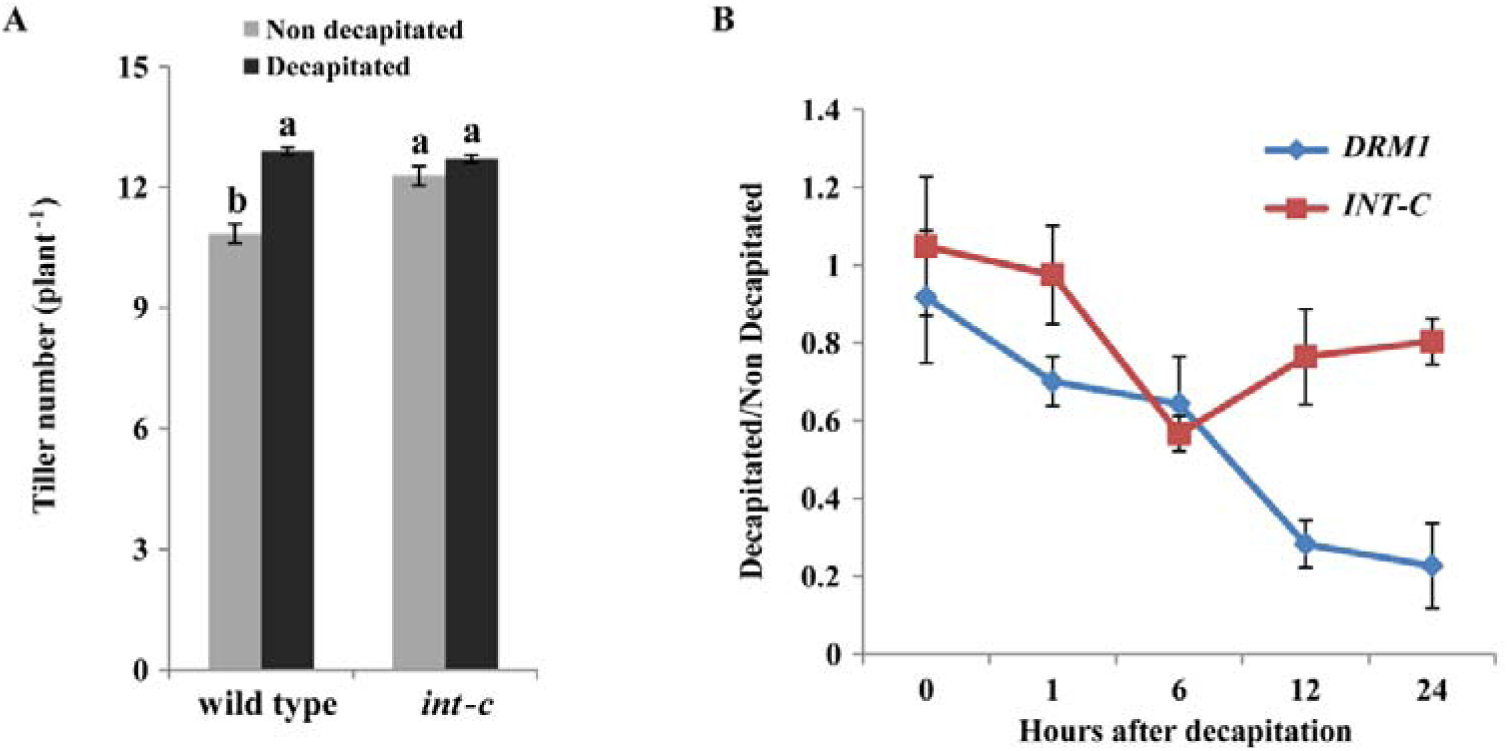
*INT-C* expression in response to decapitation. **A)** Tiller number of cv. Bowman (wild type) and *int-c* plants 2 weeks after decapitation. Bars represent means ± SD; *n* = 3 replicates with ≥16 plants. Different letters indicate significant differences according to Tukey’s test (P <0.05). **B)** Ratio of mRNA levels of *INT-C* and *DRM1* in tiller buds between decapitated and non-decapitated plants. Relative mRNA abundance of INT-C mRNA was analyzed by real-time q-RT-PCR. Bars represent means ± SD; *n* = 4 biological replicates.

*Serine/threonine protein phosphatase HvPP2A-4* was used as a reference gene. Analyzed is the early transcriptional response within 24 hours after decapitation.

### Transcriptome analysis of buds after decapitation

To investigate the transcriptome response of dormant versus activated buds, Agilent 8 × 60K customized barley microarray expression analysis (Thirulogachandar et al., 2017) using total mRNA prepared from tiller buds at the early stem elongation stage was performed. The time point 24 h after decapitation was chosen for the following two reasons: 1) down-regulation of *DRM1* expression was evident at this time point (Figure 3B) and 2) this time point was one day before the first visible effects of decapitation on the growth were detectable (2 d after decapitation, Figure 3A).

1704 differential expressed genes (DEGs) were detected in tiller buds of decapitated versus non-decapitated plants (Supplementary Table S2), among those 1011 were down-regulated and 693 up-regulated. Microarray results were confirmed by qRT-PCR for selected genes (Supplemental Figure 2). Besides *DRM1*, which was found among the down-regulated genes, several genes encoding transcription regulators were detected. 81 out of 494 DEGs regulated in opposite direction under shading conditions could be mapped to the term transcription regulator Associated with the term ABA are the auxin binding protein ABP44 and a putative ripening related bZIP Protein (CAB85632). The class of AP2-EREBP transcription regulators was found to be down regulated upon decapitation. The up-regulated genes included a large number of ribosomal proteins, cell organization, and cell cycle-related genes.

### Plant density affects *INT-C* expression

Planting density affects shoot branching in many plant species. Low plant density results in more branches, compared to growth in dense plant stands as a result of a neighbor-sensing response (Casal et al., 1986). To test the impact of plant density on *INT-C*, wild type and *int-c* plants were grown in three different densities under greenhouse conditions (1, 5 and 10 plants per pot). Tiller numbers were counted until the early stem elongation stage (Figure 4).

**Figure 4.**
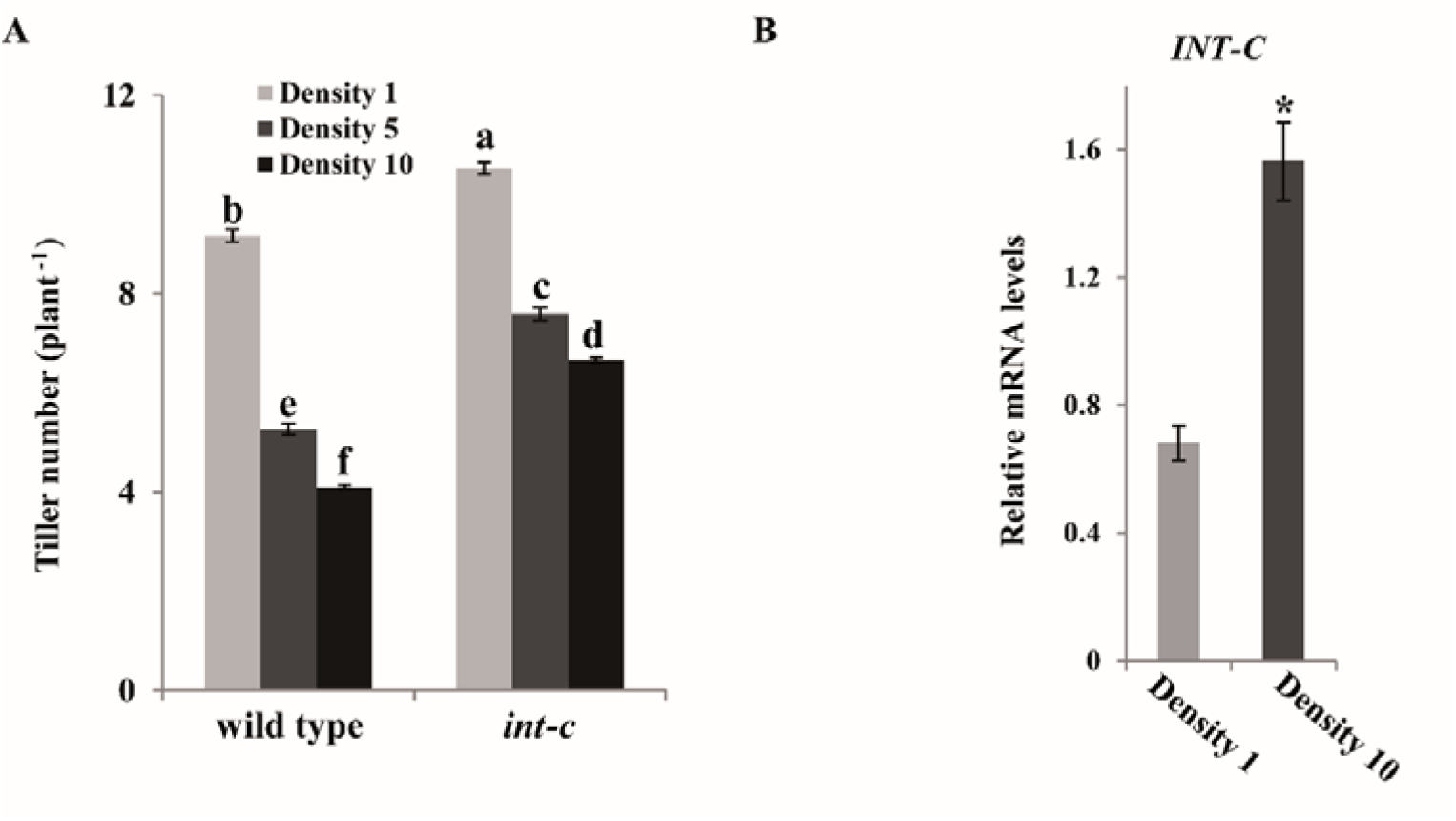
*INT-C* expression responds to planting density. **A)** Tiller number of wild type and *int-c* plants grown at planting densities of 1, 5, or 10 plants per pot. Plants were analyzed 5 weeks after sowing. Bars represent means ± SD; *n* = 3 replicates with ≥20 plants. Different letters indicate significant differences according to Tukey’s test (P < 0.01). **B)** Transcript levels of *INT-C* in the tiller bud tissue analyzed by real-time PCR at a density of 1 or 10 plants per pot. Bars represent means ± SD; *n* = 3 biological replicates. *Serine/threonine protein phosphatase HvPP2A-4* was used as a reference gene. Asterisk indicates significant difference according to Student’s *t* test at *P < 0.001.

Wild-type plants responded to increased planting density with reduced tillering. At a density of ten plants per pot, tiller bud suppression was more than half, as tiller number was 56% lower than in plants grown at one plant per pot. However, *int-c* mutants showed reduced sensitivity to this condition (33% reduction compared with plants at one plant per pot). The mRNA levels of *INT-C* were then analyzed by RT-qPCR in wild type plants grown at low (one plant per pot) and high (ten plants per pot) density. At high density, *INT-C* mRNA levels showed a two-fold increase compared to those at low density (Figure 4B). These results support the involvement of transcriptional regulation of *INT-C* on bud dormancy.

### *INT-C* is upregulated as part of the shade avoidance response

To further examine whether *INT-C* is participating in the shade avoidance response, plants were grown under two different light conditions: control light (PAR: 840 μmol m^-2^ s^-1^; R:FR ratio = 2.2), and low R:FR light mimicking shade imposition by neighboring plants (green shade, PAR: 260 μmol m^-2^ s^-1^; R:FR ratio = 0.2) (Kegge et al., 2013). To minimize a putative bias resulting from variations in leaf number and *int-c* mediated earlier flowering, plants were grown at conditions attenuating the *int-c* early flowering phenotype (Figure 5A).

**Figure 5.**
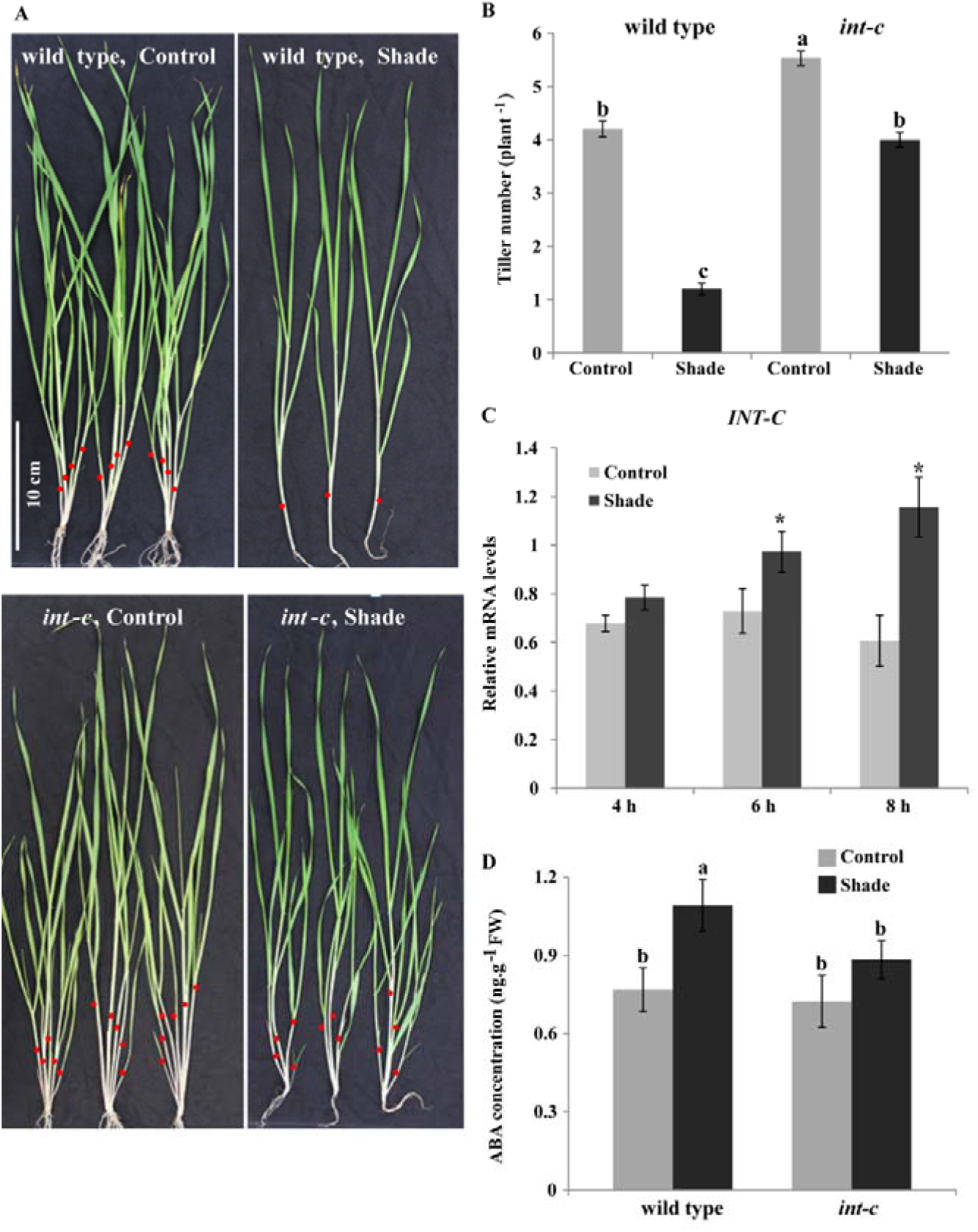
Effect of shading on tiller bud outgrowth and *INT-C* expression. **A)** Tillering phenotype of wild-type and *int-c* plants grown under control conditions or shading. Red dots indicate the primary tillers. **B)** Tiller number of wild type and *int-c* plants grown under control or shade conditions. Bars represent means ± SD; Three independent experiments with *n* ≥35 plants each. Different letters indicate significant differences according to Tukey’s test (P < 0.01). **C)** Transcript levels of *INT-C* analyzed by qPCR, in buds of shaded plants, relative to levels in control plants. Bars represent means ± SD; *n* = 3 biological replicates. *Serine/threonine protein phosphatase PP2A-4* was used as a reference gene. Asterisk indicates significant difference according to Student’s *t* test at *P < 0.05. **D)** ABA concentrations in tiller buds of wild type and *int-c* plants 6 h after exposure to shade. Bars represent means ± SD of six independent biological replicates. FW: Fresh Weight. Different letters indicate significant differences according to Tukey’s test (P < 0.05).

The shading treatment was started at the two-leaf stage. Four weeks later primary tiller number was quantified. Bowman plants grown under shade conditions responded strongly and had 4 times less tillers than plants grown in control conditions (Figure 5B), indicating that exposure of plants with young vegetative buds to a low R:FR ratio promotes bud arrest in barley. In contrast, the response of *int-c* mutants to shading was much weaker. *int-c* plants grown under shade had only 1.4 times fewer tillers than plants grown under normal light conditions (Figure 5A/B). Other phenotypic shade avoidance responses, such as hyponasty and stem elongation, were indistinguishable between the wild type and mutant plants (Figure 5A). The short-term response of *INT-C* to shade treatment was analyzed by transferring plants at the five-leaf stage (when plants had small vegetative buds) to low R:FR light. The mRNA levels of *INT-C* in tiller buds increased after 4, 6 or 8 h exposure to shade, indicating a transcriptional activation of *INT-C* in response to shade (Figure 5C). This result is in agreement with the increase of tillers in plants grown in dense stands. The higher tiller number in *int-c* coincided with a lower concentration of ABA in tiller buds (Figure 5B, D), supporting the role of ABA as mediator of INT-C-dependent tiller bud suppression in the shade.

### Transcriptome analysis of tiller buds after shading identifies *INT-C* dependent genes

To get further insight into the molecular mechanisms of *INT-C*-mediated growth responses, the transcriptome of tiller buds in wild type and *int-c* plants was analyzed. The plants were exposed to control conditions or shade (Figure 5) and the transcriptome was analyzed 6 h after exposure to shade using an Agilent 8 × 60K customized barley microarray. As *INT-C* expression was highest in buds (Figure 1), we selected this tissue for detailed analysis of RNA extraction.

While under control conditions a minor influence of *int-c-*dependent transcript changes was found, 305 differentially expressed genes (DEGs) were identified between WT vs *int-c*, out of which 185 were up- and 120 down-regulated. Under shading a stronger effect of *int-c* on the transcriptome was noted. In wild type buds, 2726 shade-responsive DEGs (1803 DEG up- and 923 DEGs down-regulated) were detected. In the *int-c* mutant, a total of 906 DEGs were identified in response to shade with 226 up-regulated genes and 680 down-regulated genes (Figure 6, Supplemental Table S2). DEGs detected by microarray were validated and confirmed for five upregulated and five downregulated genes by qRT-PCR (Supplementary Figure 3). The number of DEGs in response to shading decreased by ∼67% in *int-c* (Figure 6, Supplementary Table S2). This drop supports the hypothesis of the involvement of *INT-C* in the shade response. The overlapping DEGs, detectable in wild type and *int-c* are related to a common shade avoidance response independent of *INT-C*, while DEGs detectable in the wild type could directly or indirectly depend on *INT-C* function. These 2154 DEGs were termed *INT-C*-dependent genes of the shade response (Figure 6, Supplementary Table S2).

**Figure 6.**
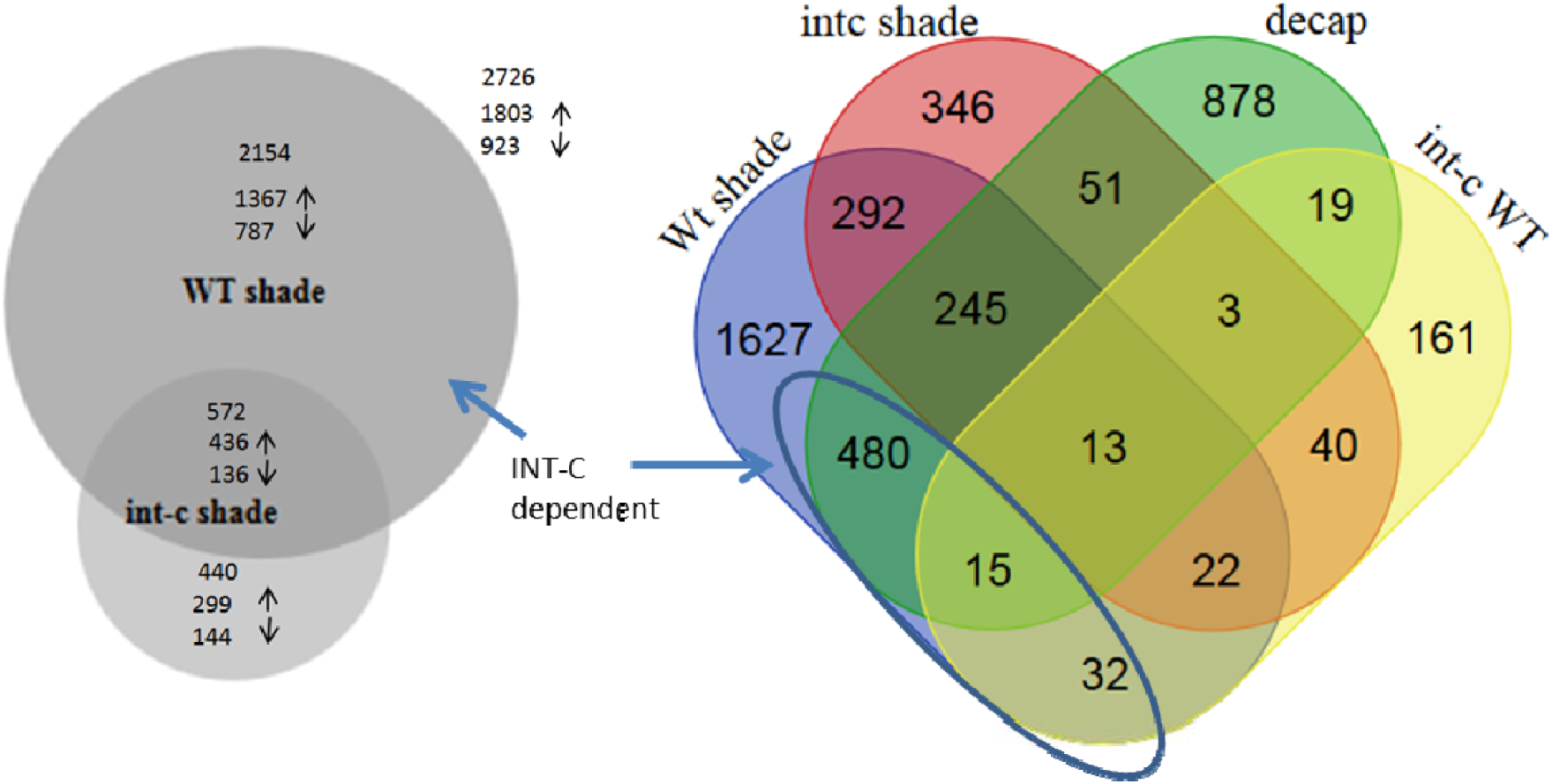
Venn diagram of differentially expressed genes (DEGs) detected after shade treatment in cv. Bowman wild type (Wt shade) and *int-c* (*int-c* shade). These blocks were compared to DEGs detectable after decapitation (decap) and *int-c* mutant and wild type (*int-c* versus WT). Numbers indicate transcript changes of fold-change ≥ 2 at an FDR of p < 0.05. (shade: INT-C mRNA induced, int-c: no functional INT-C, decap: INT-C mRNA reduced).

In Bowman buds, the upregulation of *INT-C* after shading (2.56-fold increase at an FDR of 4.78E-04 could be confirmed. In the BW421 *(int-c*.*5*) deletion mutant a similar induction of *INT-C* (2.10-fold increase, FDR = 0.0086) was detectable (Supplementary Table S2). However, in the *int-c*.*5* mutant the deletion in the *INT-C* gene leads to a frameshift in the C-terminus downstream of the R-domain. This mutation leads to a nonfunctional gene product. The inspection of the respective marker genes *HAT4/ATHB2, PIF3* and *PIF4* (Leivar and Monte, 2014) by their transcriptional activation validated the applied experimental shade conditions (Supplementary Table S2). Among the significantly upregulated genes after shade in the wild type, transcript levels of the marker gene *DRM1* that is associated with tiller bud dormancy (Stafstrom et al., 1998; Tatematsu et al., 2005) were found to be more than two-fold higher (Supplementary Table S2).

As shown in Figure 5A the shade response led to a transcriptional activation of *INT-C* and a repression of tiller outgrowth. In contrast, the decapitation of barley plants resulted in a reduced expression of *INT-C* and the opposite phenotype. As *INT-C* expression was reduced under this simulated condition, the generated dataset was used to validate the list of *INT-C*-dependent shade response genes (Supplementary Table S2).

The Venn diagram (Figure 6) illustrates 753 overlapping DEGs after decapitation and after shading (27% of 2726 DEGs after shading and 44% of 1704 DEGs after decapitation). 495 (29%) of the decapitation-induced DEGs were also detectable in response to shading. Almost all of these DEGs were oppositely regulated after decapitation and shading, respectively (Supplementary table S3). This list was defined as INT-C dependent genes.

MapMan software was used to define gene functional categories (Thimm et al., 2004; Usadel et al., 2005). For the *INT-C* dependent genes we identified hormone- and stress-related genes among the upregulated genes and cell division- and protein synthesis-related genes among the downregulated genes as the most prominent functional categories.

As all INT-C-dependent genes showed opposite responses after decapitation and under shade, these gene sets infer the causal mechanisms associated with bud activation and bud arrest, respectively. 123 genes (25 %) found to be upregulated after decapitation were downregulated under shade while 372 (75 %) genes downregulated after decapitation were upregulated under shade (Figure 6; Supplementary Table S3). During decapitation-induced bud activation, *INT-C* was rapidly downregulated (Figure 3B). Further, a strong overlap of DEGs responding to decapitation and shading was observed. Theoretically, the 372 downregulated (after decapitation) genes might be directly involved in the promotion of axillary bud arrest. This group included a number of genes related to ethylene, abscisic acid, auxin, and gibberellin signaling (AP2/ERBP, ACC, ERF1, IAA17, and GID1L2), as well as protein degradation (SKIP1, SKIP5, SKP2A, UBQ3 and UBQ4). Moreover, genes related to sugar metabolism and transport also identified within that group (TPS6, sucrose transporter). Trehalose-6-phosphate is known to be involved in sugar signaling (Figueroa and Lunn, 2016). On the other hand, the 123 upregulated genes (after decapitation) could be involved in promoting axillary bud growth. This group included many genes associated with chloroplast function and chlorophyll synthesis (chlorophyll binding protein, ATP synthase), protein synthesis (EF-Ts, ribosomal protein), and chromatin structure (HISTONE H3.2, HISTONE 2B. 3). This set of genes was co-regulated with a subset of genes related to functional categories involved in thylakoid and photosynthesis (Supplementary Figure 4).

## Discussion

The present study shows that *INT-C* is transcriptionally regulated by apical dominance and by light perception and that *INT-C* expression in tiller buds regulates tillering in response to these signals. This highlights the role of *INT-C* as a major transcriptional regulator integrating endogenous with environmental signals to determine the outgrowth of tiller buds. Based on the microarray hybridization experiment INT-C dependent genes are defined.

### *INT-C* integrates environmental signals to regulate tiller bud outgrowth

In crop plants, tillering is an important agronomic trait for yield formation. However, in barley little is known about the genetic mechanisms acting inside tiller buds to cause growth arrest. Here, we show that the bHLH transcription factor *INTERMEDIUM-C* (*INT-C*) is involved in the integration of different branching signals mediating a suppressive effect on bud outgrowth (Figure 3). INT-C itself is regulated on the transcriptional level as response of the investigated environmental conditions. This modulation of *INT-C* transcription appears to be under tight regulation of environmental and developmental stimuli that are correlated with bud outgrowth and activation of tillering. *INT-C* upregulation was observed after shading or under high planting densities (Figure 3, 4), linked to reduced tiller numbers; on the contrary suppression of *INT-C* expression, i.e., after decapitation, triggered bud outgrowth (Figure 5). This emphasizes a role of *INT-C* as a regulator integrating signals within the axillary bud to determine tiller number. This finding agrees with the results in rice where TB1 was also reported to mediate a negative function in tillering (Takeda et al., 2003; Choi et al., 2012). A comparable approach using transgenic overexpression and antisense mediated repression of INT-C resulted in the modulation of tiller and panicle development in rice. The influence and integration of environmental stimuli in rice was only addressed in response to greenhouse and paddy field conditions. In maize the involvement of the SPL gene family as direct regulators of TB1 on plant architectural traits was described and optimized plants were engineered for high-density planting (Wei et al., 2018).

### Definition of *INT-C*-dependent genes

The analysis of the shading response at the transcriptome level led to the identification of important differentially expressed genes that were directly or indirectly dependent on INT-C. Our data suggest that the number of INT-C-dependent genes (229 up- and 82 downregulated) was closely associated with tiller bud activation, confirmed through *int-c* mutant study, and INT-C function inferred from the stimulus experiments (either shade or decapitation). These target genes are promising candidates to play an important role in tiller bud transition between repression and promotion of growth. The MAPMAN-based categorization of DEGs resulted in a significant overrepresentation of the term thylakoid and genes associated with photosynthesis. In addition to this, several target genes of sink/source properties of a tissue are found (e.g. TPS6, sucrose transporters). This supports the idea of bud activation, accompanied by generation of a new initial sink tissue. Also, photosynthesis will be activated in this new formed tiller. The scheme in Figure 7 show the integration of the respective categories and important genes involved in INT-C mediated bud activation.

**Figure 7.**
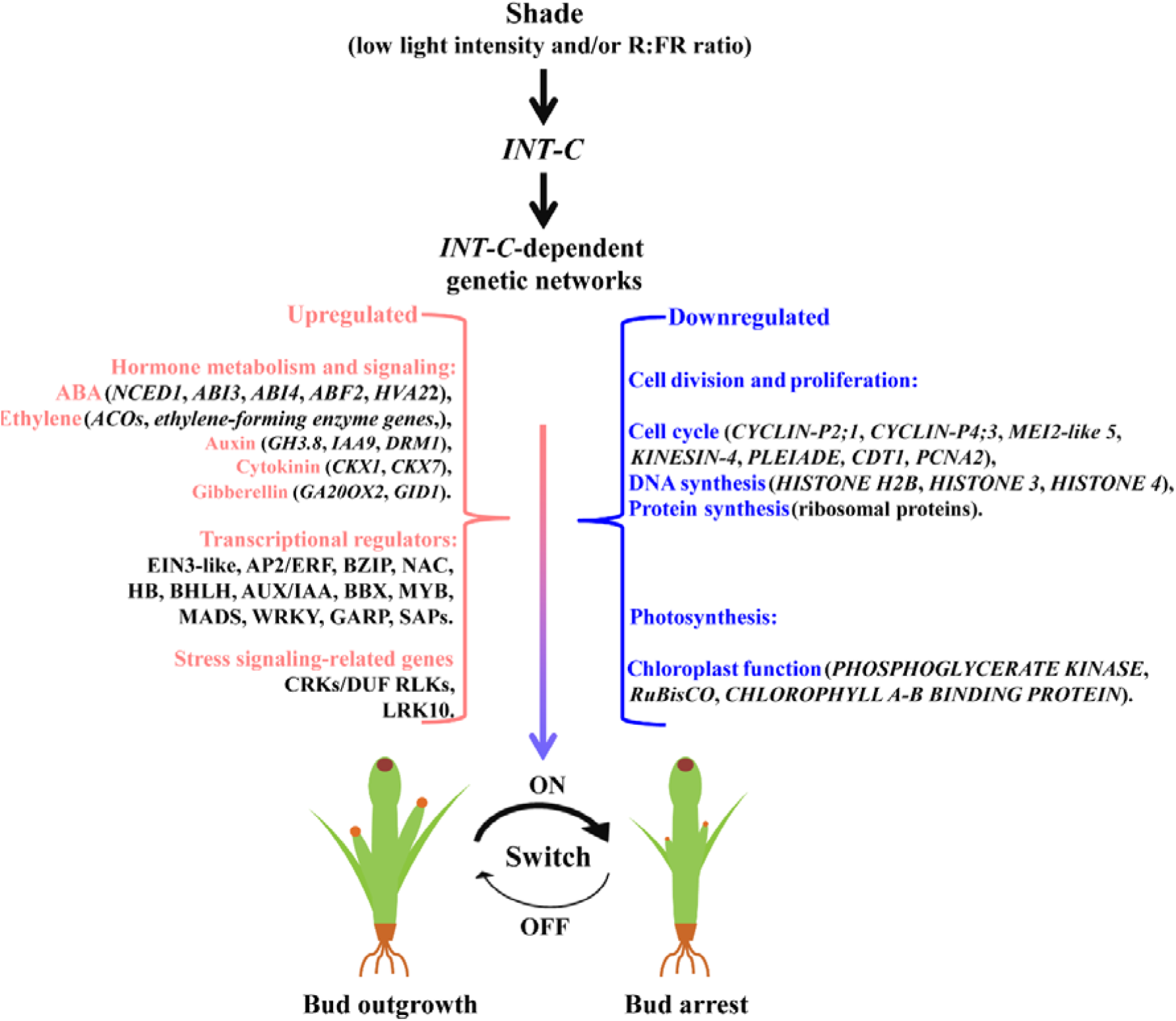
Working model for the dynamic balance of INT-C-dependent transcriptional programming to regulate tiller bud outgrowth in barley. Shade perception occurring in tiller bud activates the “master-switch” transcription factor *INT-C*, thus altering INT-C-dependent target genes, among them is a set of upregulated hormone and stress-response genes and down-regulated cell division- and ribosome-related genes. The output of this transcriptional regulation mediates bud transition from outgrowth to arrest.

### ABA signaling and the promotion of bud dormancy

ABA has been related to the maintenance and promotion of bud dormancy in many plant species: elevated ABA levels in buds are associated with the inhibition of branching during plant development (Tamas et al., 1979; Knox and Wareing, 1984; Gocal et al., 1991; Mader et al., 2003; Destefano-Beltran et al., 2006; Ruttink et al., 2007; Eggert and von Wiren, 2017), as well as in the context of responses to low R:FR ratios (Tucker and Mansfield, 1971; Reddy et al., 2013; Gonzalez-Grandio et al., 2017). Moreover, a correlation has been found between the upregulation of ABA-response genes in axillary buds and bud dormancy (Ruttink et al., 2007; Gonzalez-Grandio et al., 2013; Kebrom and Mullet, 2016). So far, neither ABA measurements have been reported in tiller buds, nor ABA-inducible genes have been studied in the tiller bud development of barley (Hussien et al., 2014). Nonetheless, it has been supposed that in barley a correlation exists between ABA signaling and bud arrest (Finkelstein, 2013). Recently, (Luo et al., 2019) and colleagues have shown that application of ABA to hydroponic cultures of rice strigolactone mutants and wild-type plants suppressed axillary bud outgrowth. Strigolactone and ABA biosynthesis share the same precursor and are closely related in regulating the tiller number in barley (Wang et al., 2018). In the present study we demonstrate that INT-C as regulatory hub regulates various ABA hormone pathway genes as well decapitation and shade responses. The large number of transcription factors found in the list of defined INT-C-dependent genes also points toward a function of *INT-C* as regulatory hub controlling a complex downstream network. Our transcript data provide evidence that wild-type tiller buds display a strong increase in the global response of ABA-related genes, while *int-c* buds, which exhibit less dormancy and continue to grow, show reduced ABA-related responses. Consistently, the number of ABA-related genes in the tiller buds was higher after shading. These findings are subsumed in the model (Figure 7) building the hypothesis that INT-C employs the ABA signaling pathway in conjunction with other hormones to mediate bud growth arrest.

## Author contributions

H.W. planned and designed the research. H.W. performed experiments and H.W., C.S. and M.K. analyzed the data. H.W., M.K., N. S, and N.v.W. conceived the study. H.W. and M.K wrote the manuscript.

## Supplemental Information

**Supplemental Figure S1**. Flowering in wild type and *int-c* mutant plants at 15 weeks after germination. Shown are three independent replicates for each.

**Supplemental Figure S2**. Validation of microarray data by qRT-PCR after decapitation treatment. A mRNA levels of a subset of genes identified as responding to the decapitation treatment both in wild-type and *int-c* samples. C and D, Pearson’s correlations between gene expression levels determined by qRT-PCR and microarray expression profiling for the same genes. Although the correlations of both datasets are high, microarray data underestimate the degree of change. The qRT-PCR and microarray data showed a very high average Pearson correlation coefficient (0,9107; confirming the high reliability of the array data.

**Supplemental Figure S3**. Validation of microarray data by qRT-PCR. A and B, mRNA levels of a subset of genes identified as responding to the shade treatment both in wild-type and *int-c* samples. C and D, Pearson’s correlations between gene expression levels determined by qRT-PCR and microarray expression profiling for the same genes. Although the correlations of both datasets are high, microarray data underestimate the degree of change. The qRT-PCR and microarray data showed a very high average Pearson correlation coefficient (0.85 and 0.81 for the wild type and *int-c*, respectively; confirming the high reliability of the array data.

**Supplemental Figure S4. Selected** MAPMAN-based categorization of DEGs

**Supplementary table 1**: List of TCP genes and associated gene expression derived from public databases.

**Supplementary table 2**: List of differential expressed genes (DEGs) for WT vs int-c (Control condition), WT vs int-c (shading condition), WT vs WT shading condition, int-c vs int-c shading condition, WT vs WT decapitation condition. Each with a list of associated direction of change.

**Supplementary table 3**: List of INT-C dependent genes, including comparison of direction after shading and decapitation.

